# Ferroptosis is executed through caspase-5 cleavage of gasdermin E in ovarian cancer cells

**DOI:** 10.64898/2026.06.15.732352

**Authors:** Mahmuda Akter, Lei Sun, Cheng Chi, Iram Hyder, Zheng Fu, Lingtao Jin, Shuang Huang

## Abstract

Ferroptosis, an intracellular iron-catalyzed form of programmed cell death (PCD) driven by lipid reactive oxygen species induced membrane damage, is mechanistically uncharacterized in its execution process. Here, we investigated ferroptosis execution in mesenchymal-like ovarian cancer cells treated with ferroptosis inducers ML162 and erastin. We showed that YVAD (a pyroptosis-associated inflammatory caspase inhibitor) and disulfiram (preventing gasdermin pore formation on plasma membrane) deterred ferroptotic cell death. Moreover, we also observed LDH release and IL-1β secretion from ferroptotic cells, suggesting that ferroptosis involves a pore-forming process. Intriguingly, ferroptosis is independent of the canonical inflammasome pathway because caspase-1 is dispensable and not activated upon ferroptosis induction. In contrast, we found that caspase-5 was activated while caspase-4 was not during ferroptosis. In addition, depletion of caspase-5 rendered cells not responding to ferroptosis inducers. Also intriguingly, GSDMD, the well-established caspase-5 substrate, was not involved in ferroptosis. We instead detected GSDME cleavage upon ferroptosis induction and knockdown of GSDME reduced cell death induced by ferroptosis inducers. As caspase-5 activity was necessary for ferroptosis and caspase-5 directly cleaved GSDME, we conclude that the axis of caspase-5/GSDME executes ferroptosis in ovarian cancer cells.

## INTRODUCTION

Ferroptosis is a form of programmed cell death (PCD) that occurs when iron-dependent lipid peroxidation overwhelms the mechanisms that normally protect polyunsaturated membrane phospholipids and maintain membrane integrity. Rather than being driven by receptor signaling, ferroptosis is regulated by interconnected metabolic pathways that include iron metabolism, membrane lipid composition, and antioxidant defense systems (Dixon et al., 2012a; X. Jiang et al., 2021; Stockwell et al., 2017). Regulation of ferroptosis depends on the delicate balance between pro-ferroptotic cellular activities that generate lipid peroxides and ferroptosis defense systems that detoxify these peroxides. In terms of pro-ferroptotic activities, the expansion of the labile iron pool provides the essential catalyst for oxidative damage while the specific phospholipid composition of the membrane also influences cellular susceptibility to ferroptosis. For example, acyl-CoA synthetase long-chain family member 4 (ACSL4) converts long chain polyunsaturated fatty acids (PUFAs) into acyl-CoA derivatives. The acyl-CoA derivatives are incorporated into membrane by lysophosphatidylcholine acyltransferase 3 (LPCAT3), and this process specifically enriches phosphatidylethanolamine (PE) species (Cui et al., 2023; Doll et al., 2017; Kagan et al., 2017a). The resulting PUFA-PEs are highly sensitive to oxidation, thus establishing a pool of peroxidation-prone substrates essential for ferroptosis. On the side of ferroptosis defense system, system X_c_⁻ (xCT), a cystine/glutamate antiporter on the plasma membrane, imports cystine into cells for glutathione (GSH) synthesis. GPX4 (glutathione peroxidase 4), a key enzyme that uses glutathione to reduce phospholipid hydroperoxides, removes destructive lipid radicals before they can kill cells (Dixon et al., 2012b; Friedmann Angeli et al., 2014). Despite major progress in defining the initiation and regulatory control of ferroptosis, the precise mechanism that translates phospholipid peroxidation to ferroptotic cell death remains unresolved.

The importance of ferroptosis was realized by the finding that the suppression of ferroptosis is essential for normal physiology. GPX4 is required for embryonic viability, and its loss *in vivo* causes catastrophic tissue injury, including acute renal failure and neurodegeneration, highlighting that proper ferroptotic control is fundamental to tissue homeostasis and cellular oxidative balance (Friedmann Angeli et al., 2014; Hambright et al., 2017). In cancer, ferroptosis is described as a natural tumor-suppressive mechanism and has thus been explored as a therapeutic vulnerability. Tumor suppressors such as p53 and BAP1 promote ferroptotic susceptibility in part by repressing SLC7A11, thereby weakening cystine-dependent antioxidant defense. Antitumor immunity can exploit the same pathway when CD8⁺ T cell-derived IFNγ suppresses cystine transport and augments lipid peroxidation in tumor cells (L. Jiang et al., 2015; W. Wang et al., 2019; Zhang et al., 2018). Recent studies suggest that Ninjurin-1 (NINJ1), a small cell-surface membrane protein, is involved in plasma membrane rupture (PMR) through its oligomerization during various lytic cell death processes including ferroptosis (Kayagaki et al., 2021; Ramos et al., 2024). However, its role appears to be both inducer and cell-type-dependent. For example, NINJ1 deficiency did not impact PMR induced by erastin while was critical for ML162-induced lysis (Bao et al., 2026; Ramos et al., 2024). Additionally, osmotic pressure, rather than ROS, triggers NINJ1 oligomerization and NINJ1 oligomerization alone does not guarantee NINJ1-dependent PMR lytic phase of ferroptosis (Dondelinger et al., 2023; Zhu et al., 2025). Together, these findings suggest that there is an unidentified executor initiating the plasma rupture for subsequent lytic process in ferroptosis.

The objective of this study is to identify the executing mechanism in ferroptosis. Since ferroptosis is recognized to have interplay with other PCD types such as necroptosis (Gong et al., 2017) and autophagy (Hou et al., 2016), we initially examined the protective effect of various specific PCD inhibitors on ML162- and erastin-induced cell death in mesenchymal-like ovarian cancer cells. Among them, only YVAD and disulfiram exhibited protection, suggesting that ferroptosis might occur in a similar mechanism of pyroptosis. Using both biochemical and genetic approaches, we demonstrate that ferroptosis necessitates caspase-5 and its activation. Furthermore, we observed the cleavage of gasdermin E (GSDME) in ferroptotic cells and showed that GSDME was essential for the occurrence of ferroptosis. Since active caspase-5 can directly cleave GSDME, we conclude that ferroptosis is executed through caspase-5 cleavage of GSDME.

## MATERIALS AND METHODS

### Cell lines and culture

Human ovarian cancer cell lines ES2, SKOV3, OVCAR8 and OVCAR3 were obtained from the American Type Culture Collection (ATCC). IGROV1 and OVCAR4 were obtained from MilliporeSigma. OCC1 was provided by Dr. Yan Xu (Indiana University). Cells were maintained in Dulbecco’s modified Eagle’s medium (DMEM) supplemented with 10% fetal bovine serum (FBS; Gibco, Cat# A4736401) at 37°C in a humidified incubator with 5% CO_2_. Cell line identity was authenticated annually by short tandem repeat profiling (PowerPlex 16 System; Johns Hopkins University Nucleic Acid Technologies, Baltimore, MD).

### Reagents and antibodies

ML162 (Cayman Chemical, Cat# 20455) and erastin (SelleckChem, Cat# S7242) were used to induce ferroptosis. Inhibitors included Z-DEVD-FMK (SelleckChem, Cat# S7312), Z-YVAD-FMK (SelleckChem, Cat# S8507), MCC950 (SelleckChem, Cat# S7809), ferrostatin-1 (SelleckChem, Cat# S7243), necrostatin-1 (Tocris, Cat# 2324/10), disulfiram (Tocris, Cat# 3807) and ADS032 (MedChemExpress, Cat# HY-156798). Primary antibodies against caspase-4 (Cat# 4450S), caspase-5 (Cat# 46680S), GSDMD (Cat# 39754S), GSDME (Cat# 88874S), and GAPDH (Cat# 2118) were purchased from Cell Signaling Technology. Antibody against ASC/PYCARD (Cat# 04-147) was obtained from EMD MilliporeSigma and antibody against caspase-1 (Cat# 22915-1-AP) was obtained from Proteintech. Antibodies against E-cadherin (Cat# sc-8426) and vimentin (Cat# sc-6260) were purchased from Santa Cruz Biotechnology.

### In vitro invasion assay

For in-vitro invasion assay, Matrigel (Corning, Cat# 356237) was diluted with DMEM at the ratio of 1:1 on ice, 100 µl of diluted Matrigel solution was then added to the upper chamber of the Transwell and incubated at 37^0^C until gel was solidified. Overnight serum-starved cells were suspended in serum-free medium and 5x10^4^ cells per well were plated on the gel in upper chamber while DMEM with 10% FBS was added in the lower chambers. After plating upper chambers into lower chambers for 48 h, contents in upper chambers were removed by cotton swab. Cells on the undersurface of upper chambers were stained with crystal violet and quantified using Image J.

### Cell viability assays and IC_50_ determination

For dose response experiments, cells were seeded at 3x10^4^ cells per well in 24-well plates overnight. Increasing concentrations of ML162 or erastin were added to cells for 24 h. Cell viability was analyzed by MTT assay. Briefly, 5mg/ml MTT (Sigma, Cat# M2128) was added to cells at final concentration of 0.5mg/ml and incubated at 37°C for 2 h. After removing medium, formazan crystals formed on the bottom of the well were solubilized in DMSO and absorbance was recorded at 570 nm using a Bio-Rad plate reader. IC_50_ values were calculated by nonlinear regression in GraphPad Prism 8.0.

### Pharmacologic inhibitor experiments

To evaluate potential involvement of known PCD mechanisms during ferroptosis, cells were seeded at 3x10^4^ cells per well in 24-well plates and pretreated for 2 h with one of the following inhibitors prior to ferroptosis induction: Z-DEVD-FMK (20µM), Z-YVAD-FMK (20µM), necrostatin-1 (1µM), chloroquine (50µM), disulfiram (1µM), MCC950 (1µM) or ADS032 (10µM). ML162 (200nM), erastin (3µM) or vehicle were then added to cells for another 24 h followed by MTT assay to determine cell viability.

### Caspase-5 activity, IL-1β secretion and LDH release assays

ES2, OCC1, and OVCAR8 cells were seeded at 1.5×10⁵ cells per well in 6-well plates and treated with ML162, erastin, or vehicle for 24 h. Cells were collected and assayed for caspase-5 activity using a colorimetric assay kit (R&D Systems, Cat# K123-100). Culture supernatants were also collected to measure IL-1β secretion by Human IL-1 beta/IL-1F2 Quantikine ELISA Kit (R&D Systems, Cat# DLB50) and LDH release by Pierce™ LDH Cytotoxicity Assay Kit (Thermo Fisher Scientific, Cat# 88953) according to manufacturers’ instructions.

### Immunoblotting

Cells were lysed in Radioimmunoprecipitation Assay buffer (50mM Tris-Cl pH 7.4, 150mM NaCl, 1% Triton X-100, 0.1% SDS, 0.5% sodium deoxycholate and 2mM EDTA) supplemented with protease inhibitor cocktail (MilliporeSigma, Cat# P8849). Protein concentration was determined using Bio-Rad Protein Assay Dye Reagent (Bio-Rad, Cat# 5000006). Lysates were boiled in loading buffer (Boster, Cat# AR1112), separated on 8–12% SDS–PAGE gels and transferred to nitrocellulose membranes. Membranes were blocked with 5% nonfat milk for 1 h and incubated with primary antibodies overnight. Membranes were then washed, incubated with HRP-conjugated secondary antibodies and signals were developed using ECL (Thermo Fisher Scientific, Cat# 32106) on a GE/Amersham Imager 680.

### siRNA-mediated knockdown

ES2, OVCAR8, and OCC1 cells were grown to 30–50% confluence in 6-well plates and then transfected with ON-TARGETplus siRNA pools (Dharmacon/Horizon Discovery) targeting CASP1 (L-004401-00-0005), CASP4 (L-004404-00-0005), CASP5 (L-004405-02-0005), GSDMD (L-016207-00-0005), GSDME (L-011844-00-0005) or PYCARD (L-004378-00-0005) using Lipofectamine 3000 (Thermo Fisher Scientific) at final siRNA concentration of 25nM. Medium was replaced after 24 h and knockdown was verified by immunoblotting with respective antibodies at 48–96 h.

### Lentiviral CRISPR–Cas9 knockout and clonal isolation

Stable CRISPR knockouts of caspase-5 were generated in OVCAR8 and OCC1 cells using lentiCRISPR v2 as described (Ran et al., 2013; Sanjana et al., 2014). Caspase-5 sgRNAs were subcloned into lentiCRISPR v2 and lentivirus was produced in HEK293T cells by co-transfection with the packaging plasmids pLP1, pLP2, and VSV-G using Lipofectamine 3000. Viral supernatants were collected and used to transduce target cells. After antibiotic selection, single-cell clones were obtained through limiting dilution. After expansion, clones were first assessed by immunoblotting using caspse-5 antibody. Clones absent of caspase-5 expression were further confirmed with genomic indel analysis at the targeted locus (Brinkman et al., 2014). Sequences of Caspase-5 sgRNA oligonucleotides are forward 5′-CACCGCCACATGCTAAAGAACAACG-3′ and reverse 5′-AAACCGTTGTTCTTTAGCATGTGGC-3′.

### GSDME cleavage assay

Full-length GSDME cDNA was subcloned into pGEXGP1. Generated constructs were transformed into BL21 competent cells and GST-Fusion proteins were purified from bacterial lysates using Glutathione Sepharose® 4B beads. Active recombinant caspase-5 was obtained from MyBioSource (Cat# MBS566581). Cleavage reactions were performed at room temperature in a total volume of 20µL using caspase buffer (50mM HEPES pH7.4, 100mM NaCl, 0.1% CHAPS, 10mM DTT and 1mM EDTA). The reaction contained 5.0µg recombinant GSDMEs and 1U of active caspase-5. Reactions were incubated at 37°C for 1 h and terminated by adding 1×Laemmli sample buffer followed by heating at 98°C for 5 min. Samples were resolved on 12% SDS–PAGE gels and stained with Coomassie prior to imaging.

### Quantification and statistical analysis

Data are reported as mean ± SEM from three independent experiments, unless stated otherwise. Statistical analysis was performed using GraphPad Prism 8.0 or Microsoft Excel. For two-group comparisons, Student’s *T*-test was used. For multiple comparisons, one-way ANOVA was applied. *P* < 0.05 was considered statistically significant.

## RESULTS

### Mesenchymal-like ovarian cancer cells are more susceptible to ferroptosis than epithelial-like ones

Previous studies show that epithelial–mesenchymal cell plasticity is closely associated with ferroptosis sensitivity in cancer, with mesenchymal-like states displaying increased susceptibility to ferroptosis. On this basis, we first verified the epithelial and mesenchymal traits in a panel of established ovarian cancer cell lines by determining their invasiveness and the abundance of E-cadherin (epithelial marker) or vimentin (mesenchymal marker). *In vitro* matrigel invasion assays showed that IGROV1, OVCAR3 and OVCAR4 cells were not invasive while OCC1, SKOV3, ES2 and OVCAR8 cells readily invaded matrigel (Fig. 1a). Parallel immunoblotting revealed E-cadherin in IGROV1, OVCAR3 and OVCAR4 cells while vimentin was enriched in OCC1, SKOV3, ES2 and OVCAR8 cells (Fig. 1b). These results established IGROV1, OVCAR3 and OVCAR4 cell lines as epithelial-like lines while OCC1, SKOV3, ES2 and OVCAR8 cell lines as mesenchymal-like ones.

**Figure 1.**
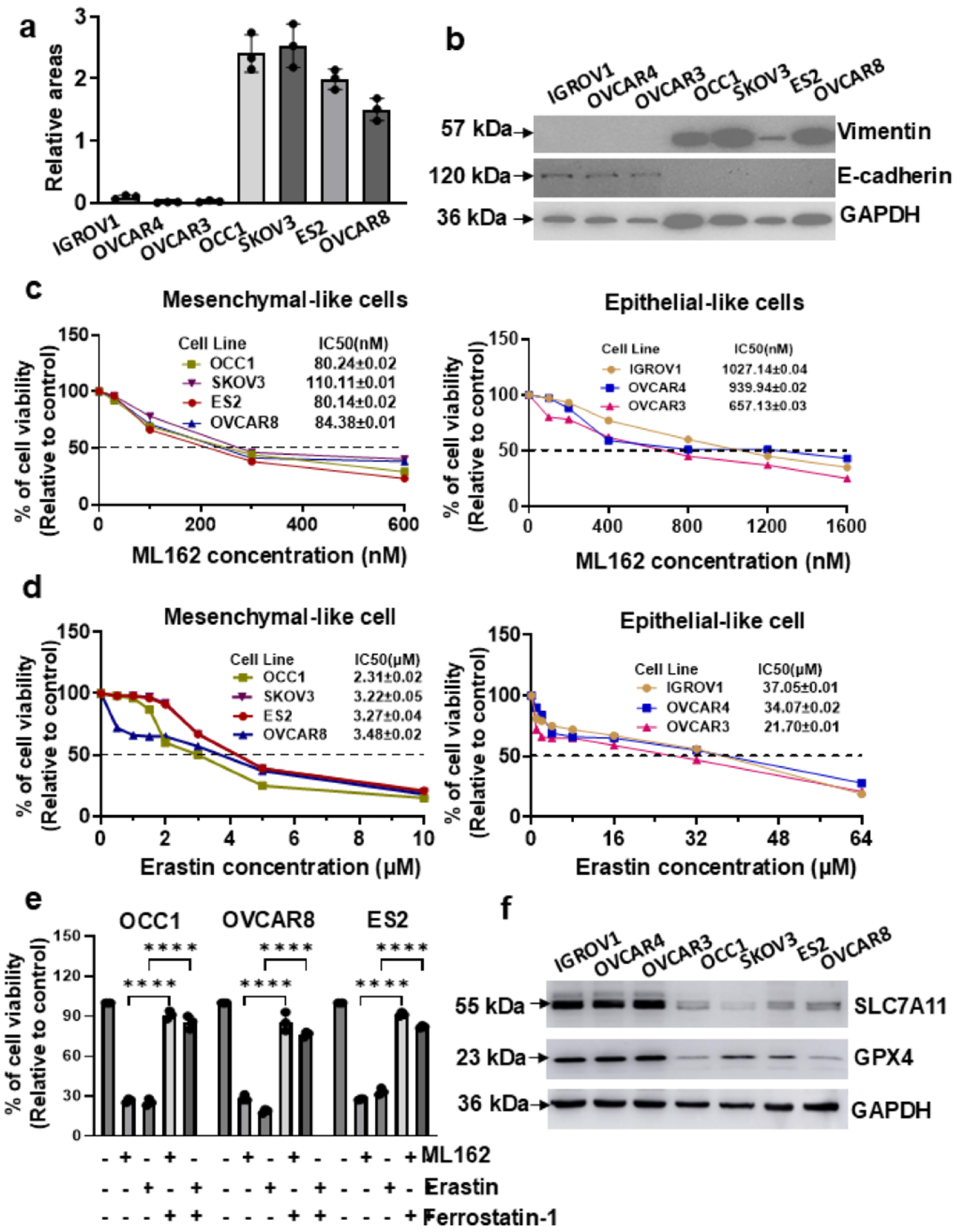
Mesenchymal-like ovarian cancer cells are more susceptible to ferroptosis than epithelial-like ones. *a*. Overnight-starved cells were plated into Matrigel-containing chambers and allowed to invade for 24 h. After removing contents in upper chambers, cells on the undersurface of chambers were stained with crystal violet and quantified using Image J. ***b***. Overnight-cultured cells were lysed and cell lysates were subjected immunoblotting with antibodies against vimentin, E-cadherin or GAPDH. ***c***. Cells were treated with varying concentration of ML162 for 24 h, followed by an MTT assay to determine cell viability. Data are means ± SEM. n = 3. ***d***. Cells were treated with varying concentration of erastin for 24 h, followed by an MTT assay to determine cell viability. Data are means ± SEM. ***e***. OCC1, OVCAR8 and ES2 cells were treated with vehicle, ML162 or erastin in the absence or presence of 10µM ferrostatin-1 for 24 h, followed by an MTT assay to analyze cell viability. % of cell viability was calculated as [(Control – Treatment)/Control] x 100. Control: vehicle-treated; Treatment: ML162 or erastin with or without ferrostatin-1. Data are means ± SEM. ****, *P* < 0.0001. ***f***. Cell lysates of overnight-cultured cells were subjected to immunoblotting with antibodies against SLC7A11, GPX4 or GAPDH.

We next examined the susceptibility of these cell lines to ferroptosis using ML162 (a GPX4 inhibitor) and erastin (an xCT inhibitor). Although ML162 and erastin reduced cell viability in all cell lines, their sensitivities were strikingly different between epithelial- and mesenchymal-like cell lines. Mesenchymal-like cell lines responded to ML162 with low-nanomolar range of IC_50_ while epithelial-like ones exhibited higher IC_50_ (Fig. 1c). Erastin produced the same overall pattern, with mesenchymal-like cell lines showing low-micromolar IC_50_ values and epithelial-like ones presenting approximately 10-fold higher IC_50_ values (Fig. 1d). To confirm that the killing caused by ML162 and erastin was truly ferroptosis, we treated OCC1, OVACR8 and ES2 cells with ferrostatin-1, a specific ferroptosis inhibitor that blocks lipid radical damage (Dixon et al., 2012; Kagan et al., 2017) followed ML162 or erastin treatment. Ferrostatin-1 pretreatment effectively blocked ML162- and erastin-induced cell death in all three cell lines (Fig. 1e).

To elucidate the mechanism underlying ferroptosis sensitivity between epithelial- and mesenchymal-like ovarian cancer cells, we examined the abundance of SLC7A11, a key subunit of xCT, and GPX4 in these cell lines. Immunoblotting showed that the levels of both SLC7A11 and GPX4 were lower in mesenchymal-like cell lines in comparison with epithelial-like ones (Fig. 1f). Further linear regression analysis revealed that IC_50_ of ML162 was positively correlated with the level of GPX4 (Supplemental Fig.S1a), suggesting that there is a negative correlation between ML162 sensitivity and GPX4 abundance. Similarly, sensitivity of erastin was also negatively correlated with the level of SLC7A11 (positive correlation between IC_50_ of erastin and SLC7A11 abundance, Supplemental Fig.S1b). These results suggest that abundance of SLC7A11 and GPX4 may be the determinant of ferroptosis sensitivity in ovarian cancer cells.

### Ferroptosis inducers efficiently kill ovarian cancer cells through pyroptosis

Because of the interplay between ferroptosis and other PCD pathways, we performed an inhibitor-based screening to determine whether any known PCD mechanisms contributed to ML162- and erastin-driven cytotoxicity. OCC1, OVCAR8 and ES2 cells were pretreated with the respective inhibitors prior to ML162 and erastin exposure. MTT assay showed that inhibiting apoptosis (DEVD), necroptosis (necrostatin-1) or autophagy (chloroquine) did not alter ML162-induced loss of cell viability (Fig. 2a). In contrast, the inflammatory caspase inhibitor YVAD drastically increased cell survival in all three lines (Fig. 2a). The same pattern was also observed with erastin (Fig. 2b). Inflammatory caspases are well characterized to trigger pyroptosis and formation of gasdermin pores is vital for pyroptotic cell death (Zou et al., 2021). We thus treated cells with disulfiram, which blocks gasdermin pore formation (C. Wang et al., 2021), followed by ML162 or erastin treatment. Pretreatment of disulfiram largely prevented cell death induced by ML162 or erastin (Fig. 2c and 2d), further supporting the notion that ferroptosis occurs in a similar mechanism of pyroptosis.

**Figure 2.**
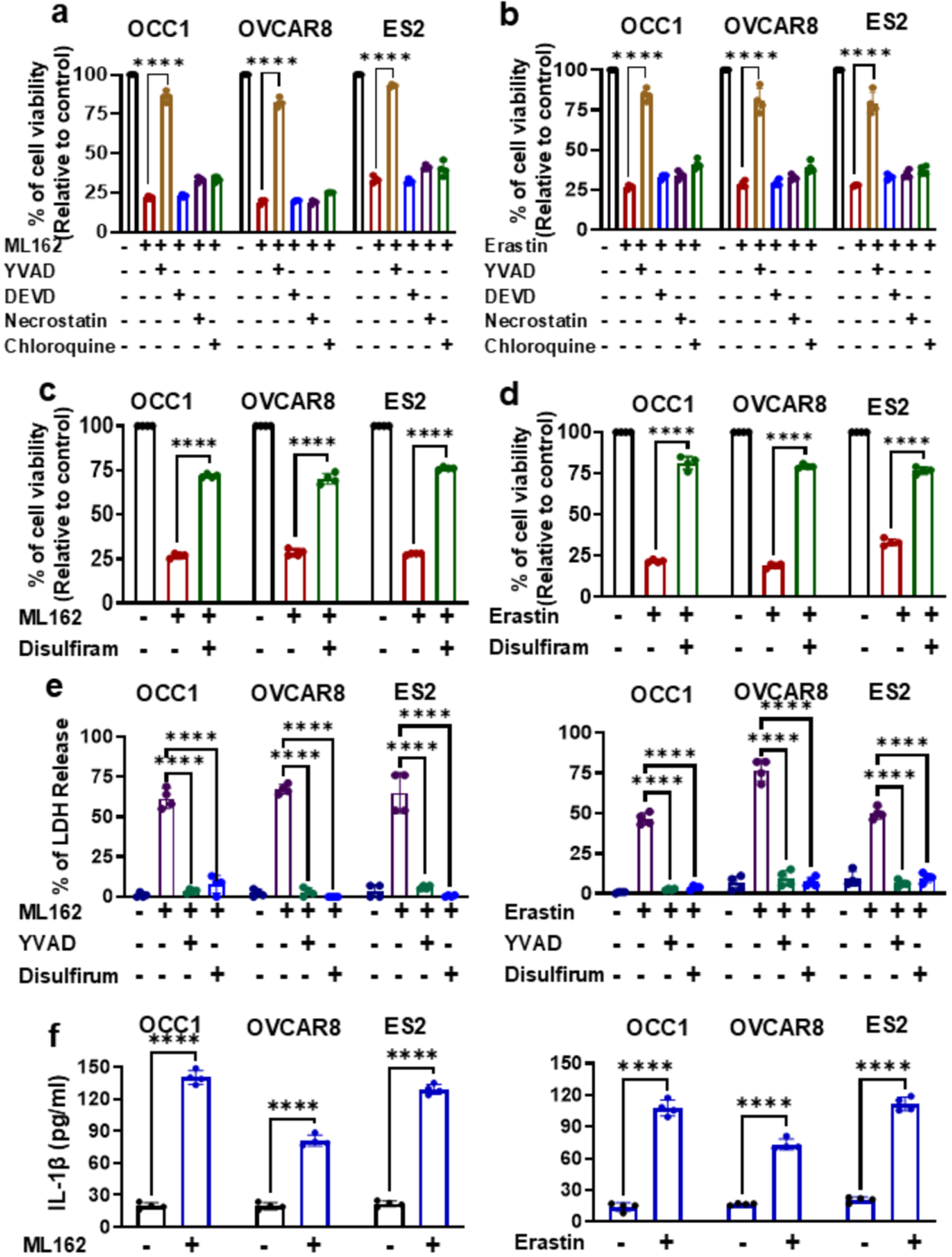
Ferroptosis occurs in a similar mechanism of pyroptosis in ovarian cancer cells. *a*. OCC1, OVCAR8 and ES2 cells were treated with vehicle or 200nM ML162 in the absence or presence of YVAD, DEVD, necrostatin or chloroquine for 24 h, followed by an MTT assay to analyze cell viability. % of cell viability was calculated as [(Control – Treatment)/Control] x 100. Control: vehicle-treated; Treatment: ML162 with or without PCD individual inhibitors. Data are means ± SEM. ****, *P* < 0.0001. ***b***. OCC1, OVCAR8 and ES2 cells were treated with vehicle or 3µM erastin in the absence or presence of individual PCD inhibitors for 24 h, followed by an MTT assay to analyze cell viability. Data are means ± SEM. ****, P < 0.0001. ***c***. OCC1, OVCAR8 and ES2 cells were treated with vehicle or 200nM ML162 in the absence or presence of disulfiram for 24 h, followed by an MTT assay to analyze cell viability. Data are means ± SEM. ****, *P* < 0.0001. ***d***. OCC1, OVCAR8 and ES2 cells were treated with vehicle or 3µM erastin in the absence or presence of disulfiram for 24 h, followed by an MTT assay to analyze cell viability. Data are means ± SEM. ****, *P* < 0.0001. ***e***. OCC1, OVCAR8 and ES2 cells were treated with vehicle, 200nM ML162 (left panel) or 3µM erastin (right panel) in the absence or presence of YVAD or disulfiram for 24 h and conditioned media were then collected to measure LDH activity. Data are means ± SEM. n = 3. ****, *P* < 0.0001. ***f***. OCC1, OVCAR8 or ES2 cells were treated with vehicle, 200nM ML162 (left panel) or 3µM erastin (right panel) for 24 h and conditioned media were then collected to measure the level of IL-1β. Data are means ± SEM. ****, *P* < 0.0001.

We next examined the morphology of OCC1 and OVCAR8 cells with or without the treatment of ML162 or erastin under a phase-contrast microscope. Both ML162- and erastin-treated cells became round and swollen, and lost membrane integrity (Supplemental Fig.S2), resembling the morphology of pyroptotic cells. In a parallel experiment, we observed that the exposure of ML162 or erastin led to marked increase in extracellular LDH release (Fig. 2e). However, LDH release led by ML162 or erastin was abolished by YVAD and disulfiram (Fig. 2e). Moreover, ML162- and erastin-treated cells secreted much higher amount of IL-1β level in supernatant compared to untreated cells (Fig. 2f). Taken together, these results suggest that ferroptotic killing is coupled to inflammatory caspase-dependent pyroptosis and that ferroptosis and pyroptosis share the similar executing mechanism.

### Ferroptosis execution is independent of canonical inflammasome pyroptosis

Pyroptosis can be driven by canonical inflammasomes (ASC-dependent caspase-1 activation) or non-canonical inflammatory caspases (caspase-4/5) (Yu et al., 2021). We initially investigated the potential role of canonical inflammasome in ferroptosis by silencing the expression of adaptor ASC/PYCARD in OCC1, OVCAR8 and ES2 cells using PYCARD siRNA pool (Supplemental Fig.S3a). MTT assay showed that knockdown of ASC did not prevent ML162 to trigger cell death in these cells (Fig. 3a). In parallel, we pretreated cells with ADS032 (NLRP1/NLRP3 inhibitor) or MCC950 (NLRP3 inhibitor) followed by ML162 exposure. Identical to what we observed with ASC knockdown, none of the inhibitors improved survival of ML162-treated cells (Fig. 3b and 3c).

**Figure 3.**
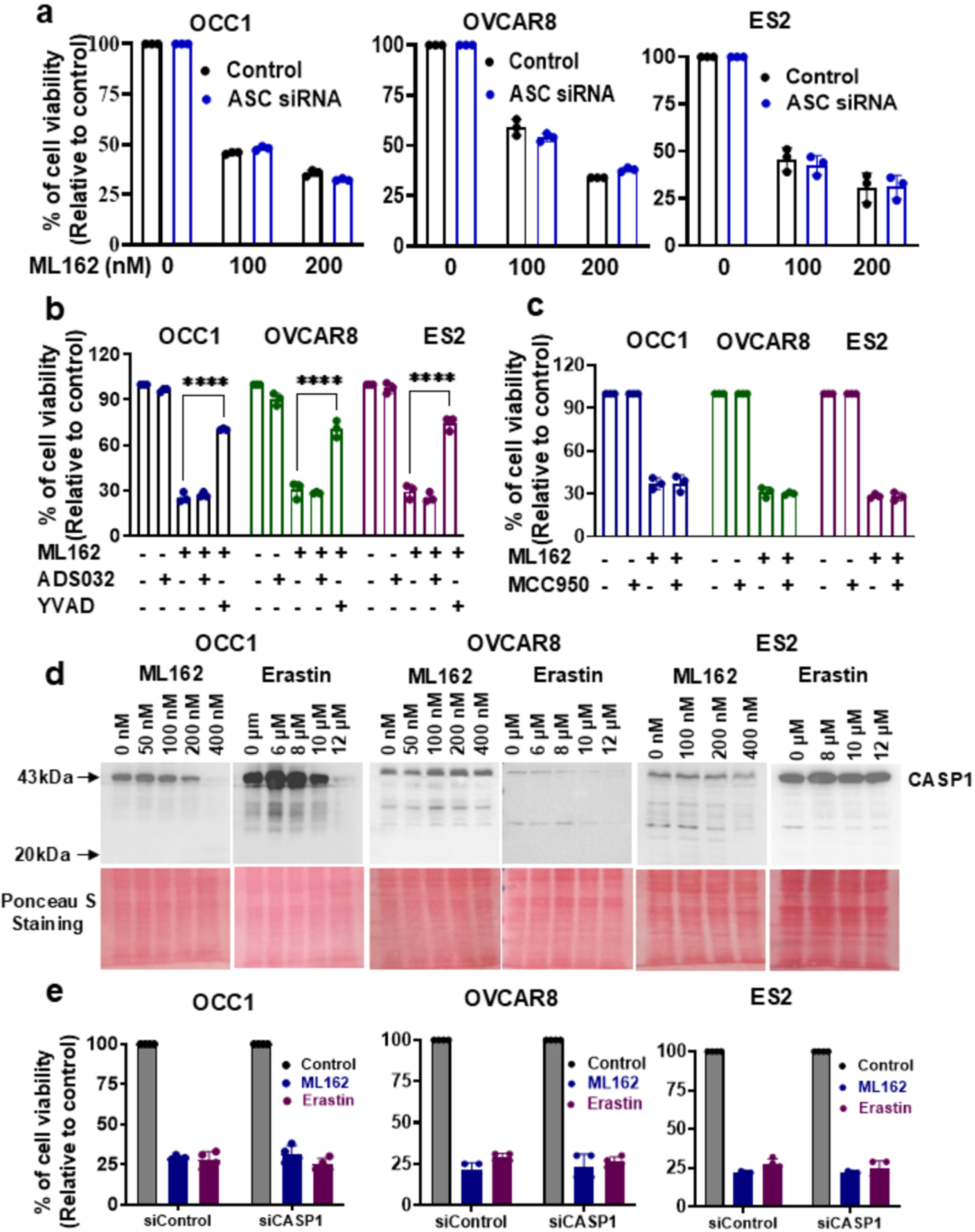
The mechanism of canonical inflammasome pyroptosis is not involved in ferroptosis. *a*. OCC1, OVCAR8 and ES2 cells were first transfected with control or ASC siRNA pool for 2 days and then treated with 200nM ML162 for 24 h, followed by an MTT assay to analyze cell viability. Data are means ± SEM. ***b***. OCC1, OVCAR8 and ES2 cells were treated with vehicle or 200nM ML162 in the absence or presence of ADS032 or YVAD for 24 h, followed by an MTT assay to analyze cell viability. Data are means ± SEM. ****, *P* < 0.0001. ***c***. OCC1, OVCAR8 and ES2 cells were treated with vehicle or 200nM ML162 in the absence or presence of MCC950 for 24 h, followed by an MTT assay to analyze cell viability. Data are means ± SEM. ***d***. OCC1, OVCAR8 and ES2 cells were treated with varying concentrations of ML162 or erastin for 24 h and then lysed for immunoblotting to detect caspase-1 (CASP1) with anti-CASP1 antibody. Membranes were stained with Ponceau S to ensure equal loading. ***e***. OCC1, OVCAR8 and ES2 cells were first transfected with control or CASP1 siRNA pool for 4 days and then treated with 200nM ML162 or 3µM erastin for 24 h, followed by an MTT assay to analyze cell viability. Data are means ± SEM.

Since caspase-1 activation can occur without ASC/inflammasome through excessive expression (Calbay et al., 2023; Padia et al., 2025), we examined the essence of caspase-1 in ferroptosis by silencing the expression of caspase-1 using CASP1 siRNA pool (Supplemental Fig.S3b). MTT assay revealed that knockdown of caspase-1 did not prevent ML162 or erastin to trigger cell death in OCC1, OVCAR8 and ES2 cells (Fig. 3d). Further immunoblotting with anti-caspase-1 antibody showed only full-length caspase-1 (FL-CASP1, 43 kDa) in all 3 cell lines across a dose range of ML162 (50–400 nM) and erastin (6–12 µM) (Fig. 3d) and the absence of cleaved caspase-1 (CL-CASP1) indicates that there is no caspase-1 activation during ferroptosis. Collectively, these results suggest that the canonical ASC/caspase-1 inflammasome pathway is not involved in ferroptosis.

### Ferroptosis necessitates caspase-5 and caspase-5 activation

Having ruled out caspase-1, we investigated the importance of non-canonical inflammatory caspases caspase-4 and 5 in ferroptosis by first examining their activation in OCC1, OVCAR8 and ES2 cells upon ML162 or erastin treatment. Immunoblotting with anti-caspase-4 antibody revealed only full-length caspase-4 (FL-CASP4, 45 kDa) in cells treated with ML162 or erastin (Fig. 4a), indicating no caspase-4 activation in ferroptotic cells. Consistently, silencing the expression of caspase-4 with CASP4 siRNA pool (Supplemental Fig.S4a) did not alter ML162- or erastin-induced loss of cell viability (Fig. 4b). In contrast, we observed that caspase-5 was processed in ferroptotic cells: amount of full-length caspase-5 (FL-CASP5, 50 kDa) decreased and cleaved caspase-5 (CL-CASP5, 40 kDa) was accumulated with the increasing concentration of ML162 or erastin in all 3 cell lines (Fig. 4a). Functionally, knockdown of caspase-5 with CASP5 siRNA pool (Supplemental Fig.S4b) substantially preserved the cell viability in all 3 cell lines exposed to ML162 or erastin (Fig. 4c). These results suggest that caspase-5 is critical for ferroptosis in ovarian cancer cells.

**Figure 4.**
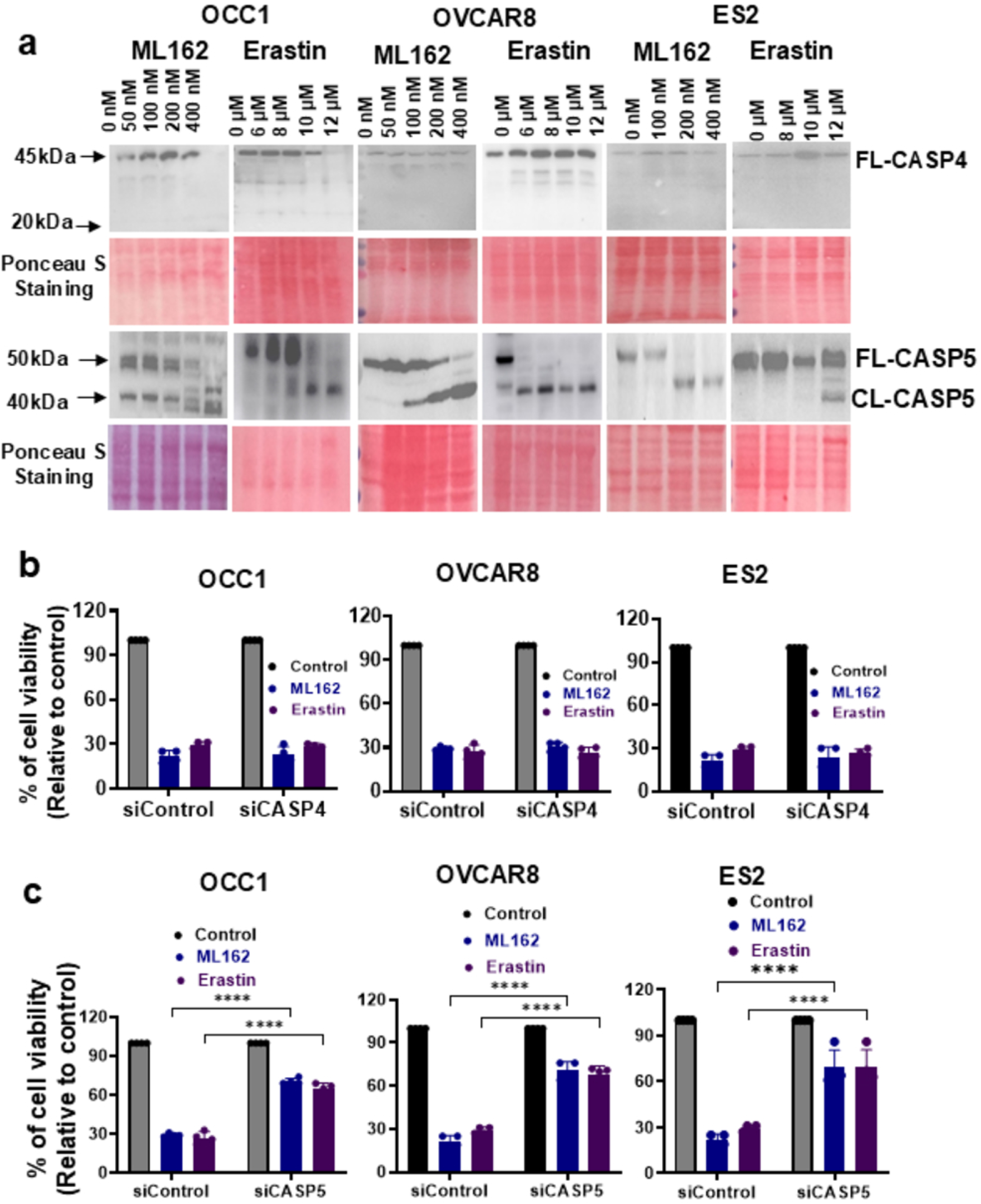
Caspase-5 is activated in cells exposed to ferroptosis inducers and its presence is necessary for ferroptosis. *a*. OCC1, OVCAR8 and ES2 cells were treated with varying concentrations of ML162 or erastin for 24 h and then lysed for immunoblotting to detect caspase-4 (CASP4) or CASP5 (CASP5) with the respective antibodies. Membranes were stained with Ponceau S to ensure equal loading. ***b***. OCC1, OVCAR8 and ES2 cells transfected with control or CASP4 siRNA pool were treated with 200nM ML162 or 3µM erastin for 24 h, followed by an MTT assay to analyze cell viability. Data are means ± SEM. ***c***. OCC1, OVCAR8 and ES2 cells transfected with control or CASP5 siRNA pool were treated with 200nM ML162 or 3µM erastin for 24 h, followed by an MTT assay to analyze cell viability. Data are means ± SEM. ****, *P* < 0.0001.

As caspase-5 may act by amplifying inflammatory signals independent of its activity (Eckhart & Fischer, 2024), we next investigated the importance of caspase-5 activation in ferroptosis by first analyzing its catalytical activity using an Ac-WEHD-pNA substrate assay. ML162 increased caspase-5 activity in both OVCAR8 and OCC1 in a dose-dependent manner (Fig. 5a). To genetically confirm the dependence of caspase-5 activity in ferroptosis, we generated CRISPR–Cas9 CASP5 knockout clones in OCC1 and OVCAR8 by initial immunoblot screening (Supplemental Fig.S5a and S5b) and subsequent DNA sequencing (Supplemental Fig.S6a and S6b). Identical to what we observed with caspase-5 knockdown, CASP5 knockout clones were resistant to ML162 treatment (Fig. 5b). We lentivirally transduced OCC1 CASP5-knockout clone 111 and OVCAR8 CASP5-knockout clone 63 with wild-type caspase-5 or enzymatically inactive caspase-5 mutant (CASP5/C328A). While the expression of wild-type caspase-5 mostly restored ML162 sensitivity, clones with the expression of catalytic inactive CASP5 mutant were still resistant to ML162 (Fig. 5c). These results suggest that the activation of caspase-5 is an obligatory step in ferroptosis.

**Figure 5.**
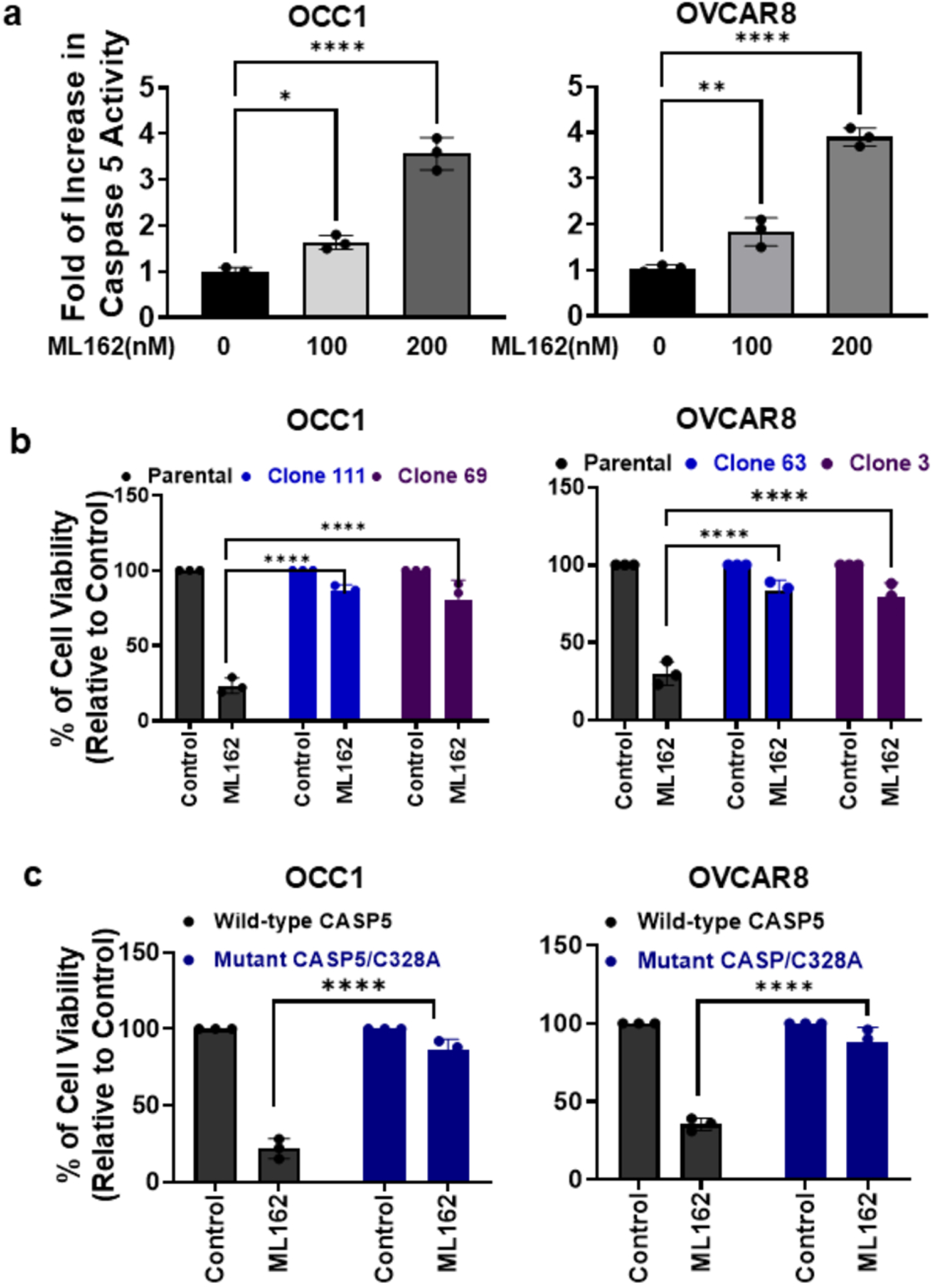
Caspase 5 activity is critical for ferroptosis. *a*. OCC1 and OVCAR8 cells were treated with varying concentrations of ML162 for 24 h followed by analyzing caspase-5 activity. Experiments were performed three times, and a representative result was shown here. Data are means ± SD. *, *P* < 0.05; **, *P* < 0.01; ****, *P* < 0.0001. ***b***. CASP5-KO clones 111 and 69 of OCC1, CASP5-KO clones 63 and 3 of OVCAR8 cells were treated with 200nM ML162 for 24 h, followed by an MTT assay to analyze cell viability. Data are means ± SEM. ****, *P* < 0.0001. ***c***. OCC1 CASP5-knockout clone 111 and OVCAR8 CASP5-knockout clone 63 were lentivirally transduced with wild-type or caspase-dead CASP5 mutant (CASP5/C328A) for 3 days and then treated with 200nM ML162, followed by an MTT assay to analyze cell viability. Data are means ± SEM. ****, *P* < 0.0001.

### Gasdermin E is involved in ferroptotic cell death

Previous studies have reported that caspase-5 carries out ferroptosis through directly cleaving GSDMD (Eckhart & Fischer, 2024) and we also observed that disulfiram blocked ferroptosis (Fig. 2b). Because disulfiram was described to primarily target GSDMD (Hu et al., 2020), we asked whether GSDMD partook ferroptosis in ovarian cancer cells. Immunoblotting showed that treatment of ML162 or erastin did not result in GSDMD cleavage (Supplemental Fig.S7). Consistently, knockdown of GSDMD with GSDMD siRNA pool (Supplemental Fig.S8a) exhibited little effect on ML162- or erastin-triggered cell death (Fig.S8b), thus ruling out the participation of GSDMD in ferroptosis.

Besides GSDMD, disulfiram and its active metabolite CuET have been shown to inhibit GDDME cleavage (C. Wang et al., 2021). In fact, exposure of OCC1, OVACR8 and ES2 cells to ML-162 or erastin all led to GSDME processing - reduced amount of full-length GSDME (FL-GSDME, ∼50 kDa) and accumulated level of cleaved fragment (CL-GSDME, ∼35 kDa) in a dose-responsive manner (Fig. 6a). Functionally, silencing the expression of GSDME with GSDME siRNA pool (Supplemental Fig.S9) drastically improved the cell viability in cells under the treatment of ML162 or erastin (Fig. 6b). These results establish GSDME as the pore-forming effector required for ferroptosis in ovarian cancer cells.

**Figure 6.**
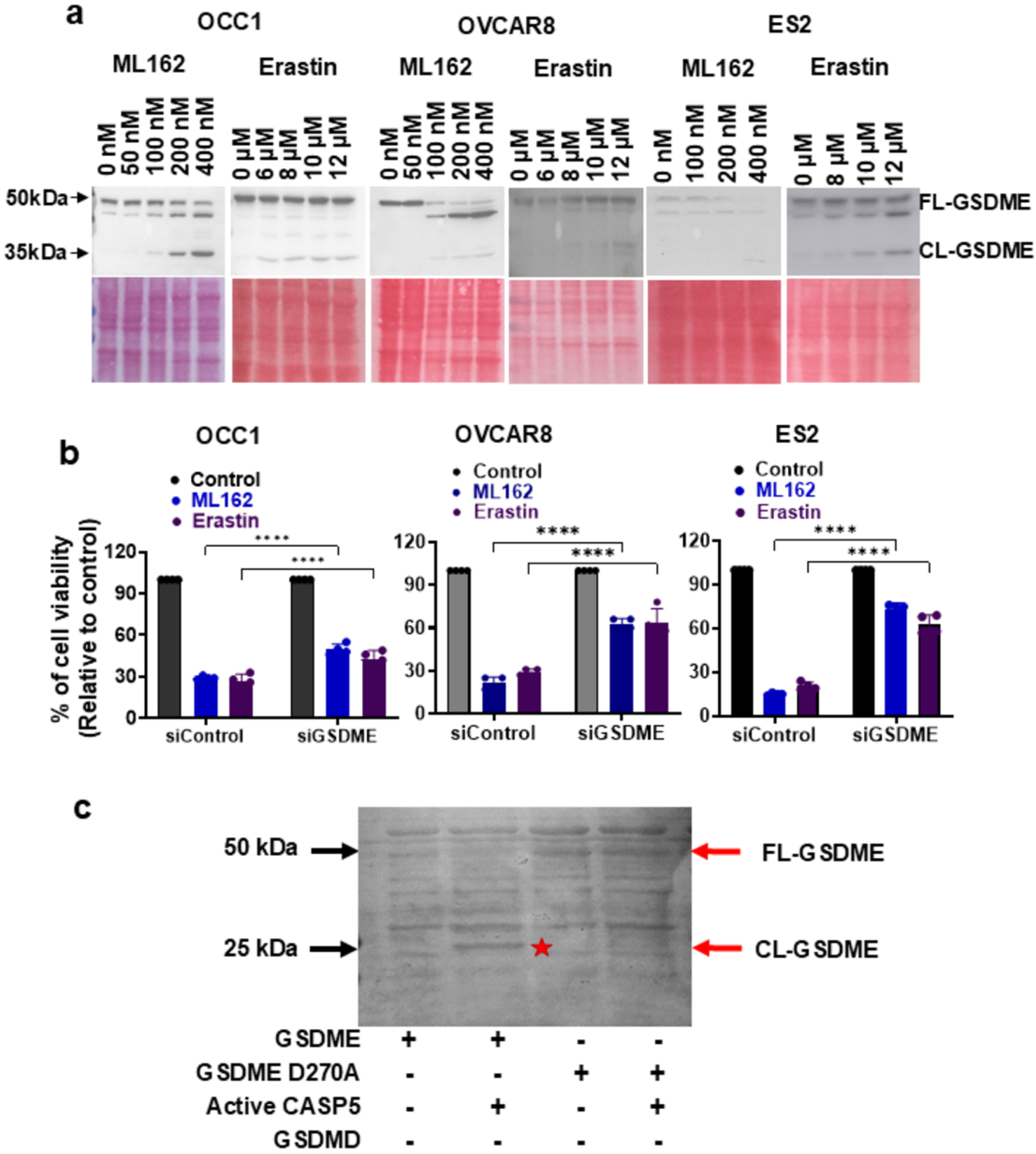
GSDME is cleaved by caspase-5 to be engaged in ferroptosis. *a*. OCC1, OVCAR8 and ES2 cells were treated with varying concentrations of ML162 or erastin for 24 h and then lysed for immunoblotting to detect GSDME with the respective antibodies. Membranes were stained with Ponceau S to ensure equal loading. ***b***. OCC1, OVCAR8 and ES2 cells transfected with control or GSDME siRNA pool were treated with 200nM ML162 or 3µM erastin for 24 h, followed by an MTT assay to analyze cell viability. Data are means ± SEM. ****, *P* < 0.0001. ***c***. 5.0µg of recombinant wild-type GSDME or GSDME mutant (D270A) was incubated with 1U of active caspase-5 in caspase reaction buffer at 37°C for 1 h. After termination the reactions, samples were resolved on 12% SDS–PAGE gels, stained with Coomassie and imaged. Red star denotes the band of cleaved GSDME (CL-GSDME). Red arrow points the position of full-length GSDME (FL-GSDME).

Recent studies have shown that GSDME is cleaved by apoptotic caspase-3 (Rogers et al., 2017). To determine whether ML162 activated caspase-3, we examined caspase-3 activation in ML162-treated OCC1 and OVCAR8 cells and were unable to detect active caspase-3 (CL-CASP3, 17 kDa) (Supplemental Fig.S10). These results are coherent with the observation that DEVD did not affect ferroptosis in our inhibitor screen experiment (Fig. 2a). To determine whether caspase-5 can directly cleave GSDME, we generated recombinant wild-type GSDME and GSDME mutant carrying cleavage site mutation (GSDME/D270A). Reconstituted cleavage experiment showed that active recombinant caspase-5 cleaved wild-type GSDME but not GSDME/D270A (Fig. 6d). Together, these results suggest that caspase-5 performs ferroptosis by directly cleaving GSDME to generate pore-forming effector critical for lytic cell death.

## DISCUSSION

Ferroptosis is commonly described as an iron-dependent cell death program driven by overwhelming lipid peroxidation. Iron contributes to lipid peroxidation, while cells rely on the cystine–glutathione–GPX4 system to prevent this damage from becoming lethal. When these defenses fail, toxic lipid peroxides build up and membranes lose their normal structure and function (Dixon et al., 2012a; Doll et al., 2017; Kagan et al., 2017a; Stockwell, 2022). Although ferroptosis has been well characterized at the level of initiation and control, the molecular basis of the downstream execution mechanism remains unclear. This study intends to fill this knowledge gap by identifying the executor of ferroptosis. Using mesenchymal-like ovarian cancer cells, we demonstrate that cell death elicited by ferroptosis inducers ML162 and erastin was effectively blocked by inflammatory caspase inhibitor YVAD and disulfiram. Moreover, we detected marked increase in LDH release and IL-1β secretion from ferroptotic cells. Since pyroptosis is characterized by LDH release, IL-1β secretion and being inhibited by disulfiram, these results suggest that ovarian cancer cells undergo ferroptosis in a mechanism resembling pyroptosis. In fact, final stage of ferroptosis displays cell swelling and membrane blebbing, which is typical for lytic cell death like pyroptosis. Also similar to pyroptosis, recent studies show that ferroptotic cells emit various damage-associated molecular pattern (DAMP) molecules such as HMGB-1, ATP and DNA (Efimova et al., 2020; Wen et al., 2019; Wiernicki et al., 2022).

Pyroptosis can undertake either canonical or non-canonical inflammasome pathway, in which the former hinges on caspase-1 while the later depends on caspase-4/5 in humans and caspase-11 in mice. Although YVAD can inhibit all inflammatory caspases, it is regarded as more selective to caspase-1. We thus initially hypothesized that ferroptosis occurs in a caspase-1/inflammasome dependent mechanism in ovarian cancer cells. However, we failed to detect caspase-1 activation and silencing caspase-1 did not affect cell death induced by ML162 or erastin. Moreover, knockdown of ASC, inhibiting NLRP1 or NLRP3 with selective inhibitors exhibited negligible effect on ML162-induced cell death. Together, these results indicate that ferroptosis does not occur in the canonic inflammasome pathway. In contrast, we observed caspase-5, but not caspase-4 activation during ferroptosis in ovarian cancer cells. Consistently, depleting caspase-5, but not caspase-4 rendered cells resistant to ferroptosis inducers, supporting the notion that ferroptosis ensues in a non-canonical inflammasome pathway involving in caspase-5 in ovarian cancer cells. Caspase-5 is primarily activated by cytosolic lipopolysaccharide (LPS) during Gram-negative bacterial infection (Viganò et al., 2015). Heme was also reported to activate caspase-5 independently of canonical inflammasome (Bolívar et al., 2021). Our findings demonstrate that ferroptosis inducers activate caspase-5 and caspase-5 activation is critical for ferroptosis, providing evidence that ferroptosis inducers are functional caspase-5 activators. Activation of caspase-5 by LPS requires its direct binding to lipid A moiety of LPS (Shi et al., 2014). As inflammatory caspases can also bind oxidized 1-palmitoyl-2arachidonoyl-sn-glycero-3-phoorylcholine (a type of oxidized phospholipid) (Chu et al., 2018; Kagan et al., 2017b; Zanoni et al., 2016), we consider the possibility of the particular peroxidized phospholipids produced in the early phase of ferroptosis as the activator of caspase-5 through its direct interaction with caspase-5.

Gasdermins are a family of pore-forming proteins that serve as key executors of pyroptosis (Shi et al., 2017). Proteolytic cleavage of gasdermins releases N-terminal fragments that oligomerize in membranes and form pores, leading to ionic imbalance, swelling, and rupture (Ding et al., 2016; Shi et al., 2015). Among all gasdermins, GSDMD is best described and is the substrate of all inflammatory caspases including caspase-5 (Shi et al., 2015). In this study, we instead found that GSDME is the one responsible for ferroptosis in ovarian cancer cells. Prior works reported that overexpression of caspase-5 in HEK293 cells resulted in caspase-3 activation (Eckhart et al., 2006; Eckhart & Fischer, 2024) and that caspase-3 can cleave GSDME in normal epithelial cells (Rogers et al., 2017). However, we failed to detect caspase-3 activation and caspase-3 inhibitor DEVD was ineffective to deter cell death elicited by ML162 or erastin. These findings suggest that direct cleavage of GSDME by caspase-5 results in ferroptosis in ovarian cancer cells. This notion is further supported by our observation that activate caspase-5 was able to cleave recombinant wild-type GSDME but not the GSDME carrying caspase-resistant mutation.

Epithelial–mesenchymal cell plasticity is closely associated with ferroptosis sensitivity in cancer, with mesenchymal-like states showing increased susceptibility (Schwab et al., 2024). We similarly observed that mesenchymal-like ovarian cancer cell lines are 10-fold more sensitive to ML162 or erastin than epithelial-like ones. This feature is very helpful for ovarian cancer therapy because platinum-based therapy is the cornerstone treatment in ovarian cancer and mesenchymal-like ovarian cancer is generally much more resistant to platinum drugs (Loret et al., 2019; Marchini et al., 2013). Early studies attributed higher ferroptosis sensitivity in mesenchymal-like cells to lipid metabolic reprogramming and phospholipid composition changes. For example, zinc finger E-box-binding homeobox-1 shifts the expression of key lipogenic enzymes including stearoyl-CoA desaturase-1, fatty acid synthase, fatty acid desaturase-2, elongation of very long-chain fatty acid protein-5 and acyl-CoA synthetase long-chain family member-4, which increase the proportion of pro-ferroptotic polyunsaturated fatty acid-containing phospholipids, thus a membrane environment more permissive to lipid peroxidation and ferroptotic damage (Lee et al., 2020; Schwab et al., 2024). In our study, we observed that the levels of GPX4 and SLC7A11 are lower in mesenchymal-like ovarian cancer cells than epithelial-like ones. Our results suggest that, in addition to enhanced pro-ferroptotic cellular activities that generate lipid peroxides, decreased levels of ferroptosis-defensive proteins such as SLC7A11 and GPX4 may also play a critical role in ferroptosis susceptibility because lower abundance of these proteins renders mesenchymal-like cells with poorer capacity to tackle lipid peroxidation and thus more sensitive ferroptosis induction.

Our findings that ferroptosis occurs in a mechanism of pyroptosis hold a tremendous promise to ovarian cancer immunotherapy because ovarian cancer is unresponsive to immunotherapy. A recent study showed that induction of pyroptosis by nanoparticle-delivered active GSDMD greatly sensitized mammary tumors to anti-PD1 therapy (Q. Wang et al., 2020). In this study, pyroptosis of less than 15% of tumor cells in a tumor was sufficient to provoke robust therapeutic effect of checkpoint inhibitors. Because of the nature of ferroptosis as pyroptosis, we reckon that agents triggering ferroptosis will bestow non-immunogenic ovarian cancer responding to immunotherapy. In future study, we will investigate the potential of such therapeutic strategy in ovarian cancer models.

In summary, our study has addressed the knowledge gap in ferroptosis execution. We demonstrate that caspase-5 cleavage of GSDME is the terminal execution step in ovarian cancer cells (may also include other cell types). We believe that our findings will provide new opportunities to manipulate ferroptotic sensitivity for therapeutic benefit.

## ACKNOWLEDGEMENT

This is part of the doctoral work of Mahmuda Akter at the University of Florida College of Medicine.

## Data and materials availability

All data needed to evaluate the conclusions in the paper are present in the paper and/or the Supplementary Materials.

